# Panoramic stitching of heterogeneous single-cell transcriptomic data

**DOI:** 10.1101/371179

**Authors:** Brian Hie, Bryan Bryson, Bonnie Berger

## Abstract

Researchers are generating single-cell RNA sequencing (scRNA-seq) profiles of diverse biological systems^1–4^ and every cell type in the human body.^5^ Leveraging this data to gain unprecedented insight into biology and disease will require assembling heterogeneous cell populations across multiple experiments, laboratories, and technologies. Although methods for scRNA-seq data integration exist^6,7^, they often naively merge data sets together even when the data sets have no cell types in common, leading to results that do not correspond to real biological patterns. Here we present Scanorama, inspired by algorithms for panorama stitching, that overcomes the limitations of existing methods to enable accurate, heterogeneous scRNA-seq data set integration. Our strategy identifies and merges the shared cell types among all pairs of data sets and is orders of magnitude faster than existing techniques. We use Scanorama to combine 105,476 cells from 26 diverse scRNA-seq experiments across 9 different technologies into a single comprehensive reference, demonstrating how Scanorama can be used to obtain a more complete picture of cellular function across a wide range of scRNA-seq experiments.

## Main Text

Individual single-cell RNA sequencing (scRNA-seq) experiments have already been used to discover novel cell states^1,2^ and reconstruct cellular differentiation trajectories^3,4^. Through global efforts like the Human Cell Atlas^5^, researchers are now generating large, comprehensive collections of scRNA-seq data sets that profile a diversity of cellular function, which promises to make a substantial contribution to our understanding of fundamental biology and disease. Assembling a large, unified reference data set, however, may be compromised by differences due to experimental batch, sample donor, or sequencing technology. While recent approaches have shown that it is possible to integrate scRNA-seq studies across multiple experiments^6,7^, these approaches automatically assume that all data sets share at least one cell type in common^6^ or that the gene expression profiles share largely the same correlation structure across all data sets.^7^ These methods therefore have limited utility when integrating collections of data sets with considerable differences in cellular composition such as the Human Cell Atlas.

Here we present Scanorama, a strategy for efficiently integrating multiple scRNA-seq data sets even when they are composed of heterogeneous cellular phenotypes. Scanorama takes inspiration from computer vision algorithms for panorama stitching that identify images with overlapping content, which are then merged into a larger panorama (**Fig. 1a**).^8^ Analogously, Scanorama automatically identifies only those scRNA-seq data sets with overlapping cell types and can leverage those overlaps for batch-correction and integration (**Fig. 1b**), without also merging data sets that do not overlap. Importantly, Scanorama is robust to different data set sizes and sources, preserves data set-specific populations, and does not require that all data sets share at least one cell population, e.g., using a cell control spiked into all data sets.^6^

**Figure 1.**
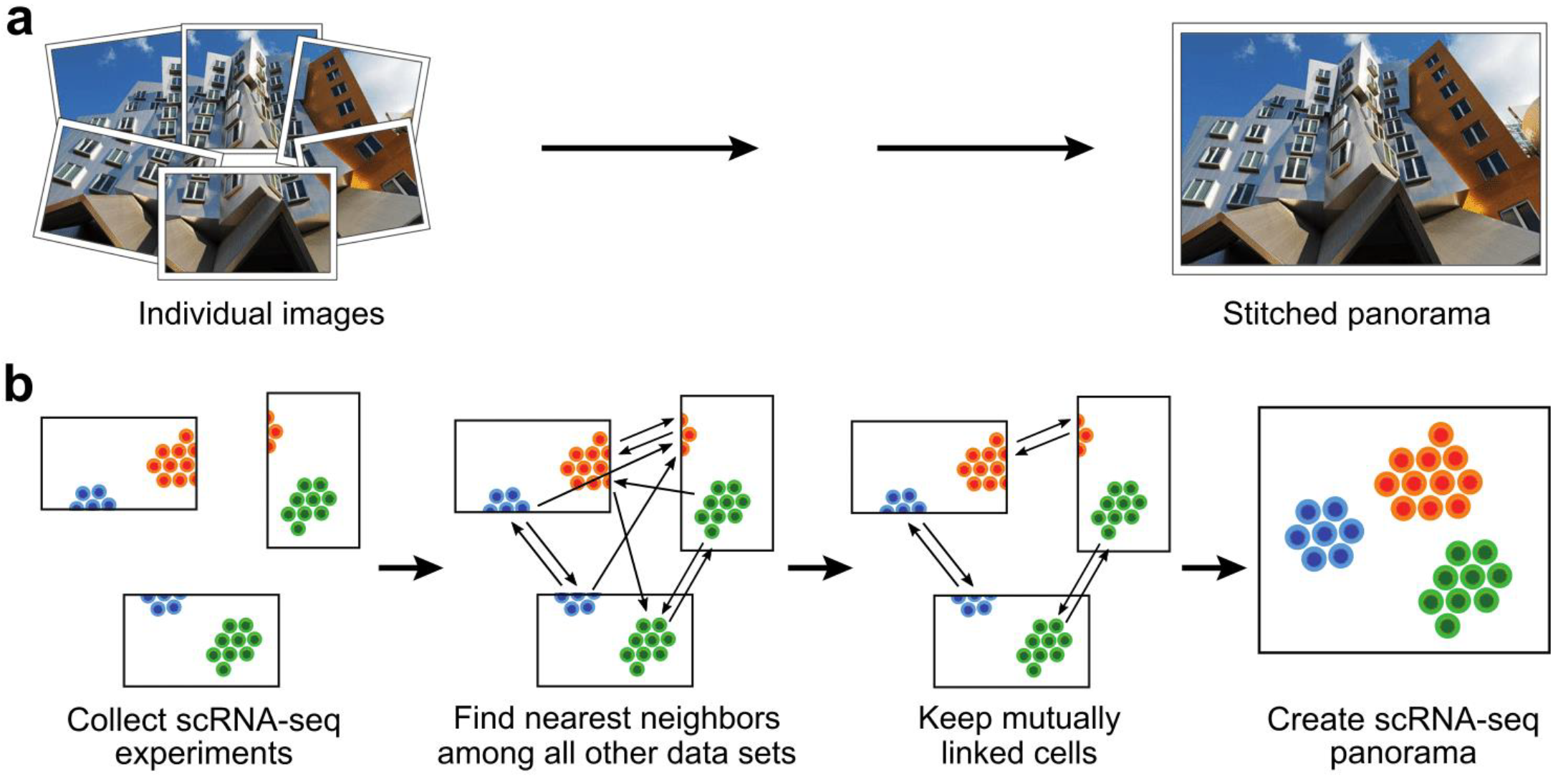
Illustration of “panoramic” data set integration. (**a**) A panorama stitching algorithm finds and merges overlapping images to create a larger, combined image. (**b**) A similar strategy can also be used to merge heterogeneous scRNA-seq data sets. Scanorama searches nearest neighbors to identify shared cell types among *all pairs of data sets*. Dimensionality reduction techniques and an approximate nearest neighbors algorithm based on hyperplane locality sensitive hashing and random projection trees greatly accelerate the search step. Mutually linked cells form matches that can be leveraged to correct for batch effects and merge experiments together (**Methods**).

Our approach generalizes mutual nearest neighbors matching, a technique which finds similar elements between two data sets, to instead find similar elements among a large number of data sets. Originally developed for pattern matching in images^9^, finding mutual nearest neighbors has also been used to identify common cell types between two scRNA-seq data sets at a time.^6^ However, to align more than two data sets, existing methods^6,7^ select one data set as a reference and successively integrate all other data sets into the reference one at a time, which may lead to suboptimal results depending on the order in which the data sets are considered (**Supplementary Fig. 1**). In contrast, Scanorama is not vulnerable to this problem since it finds matches between *all pairs of data sets*. Conceptually, we first form directed links from cells in one data set to their nearest neighbors among all other data sets, where cells with more similar gene expression profiles are also closer together, which we repeat for each data set. We then match cells between all pairs of data sets only if they were mutually linked in the previous search step, which increases the robustness of our matches and excludes links involving data set-specific populations. Once cell type matches have been determined, they can then be used to merge data sets together to create scRNA-seq “panoramas” (**Methods**).

Although searching for matching cells among all data sets may seem computationally intractable, we introduce two key optimizations to greatly accelerate our matching procedure. Instead of performing the nearest neighbor search in the high-dimensional gene space, we compress the gene expression profiles of each cell into a low-dimensional embedding using an efficient, randomized singular value decomposition (SVD)^10^ of the cell-by-gene expression matrix. Additionally, we use an approximate nearest neighbor search based on hyperplane locality sensitive hashing^11^ and random projection trees^12^ to greatly reduce the nearest neighbor query time both asymptotically and in practice (**Methods**).

To verify the merit of our approach, we first tested Scanorama on simulated data and a small collection of scRNA-seq data sets. We simulated three data sets with four cell types in total but where the first and third data sets had no cell types in common (**Supplementary Fig. 2a-c**). We also obtained three previously generated^13^ data sets: one of 293T cells, one of Jurkat cells, and one with a 50:50 mixture of 293T and Jurkat cells (**Fig. 2a**). In both cases, we were able to merge common cell types across data sets (**Fig. 2b, Supplementary Fig. 2d**) without also merging disparate cell types together, as was the case when applying existing integration methods to these data sets (**Fig. 2c,d**).

**Figure 2.**
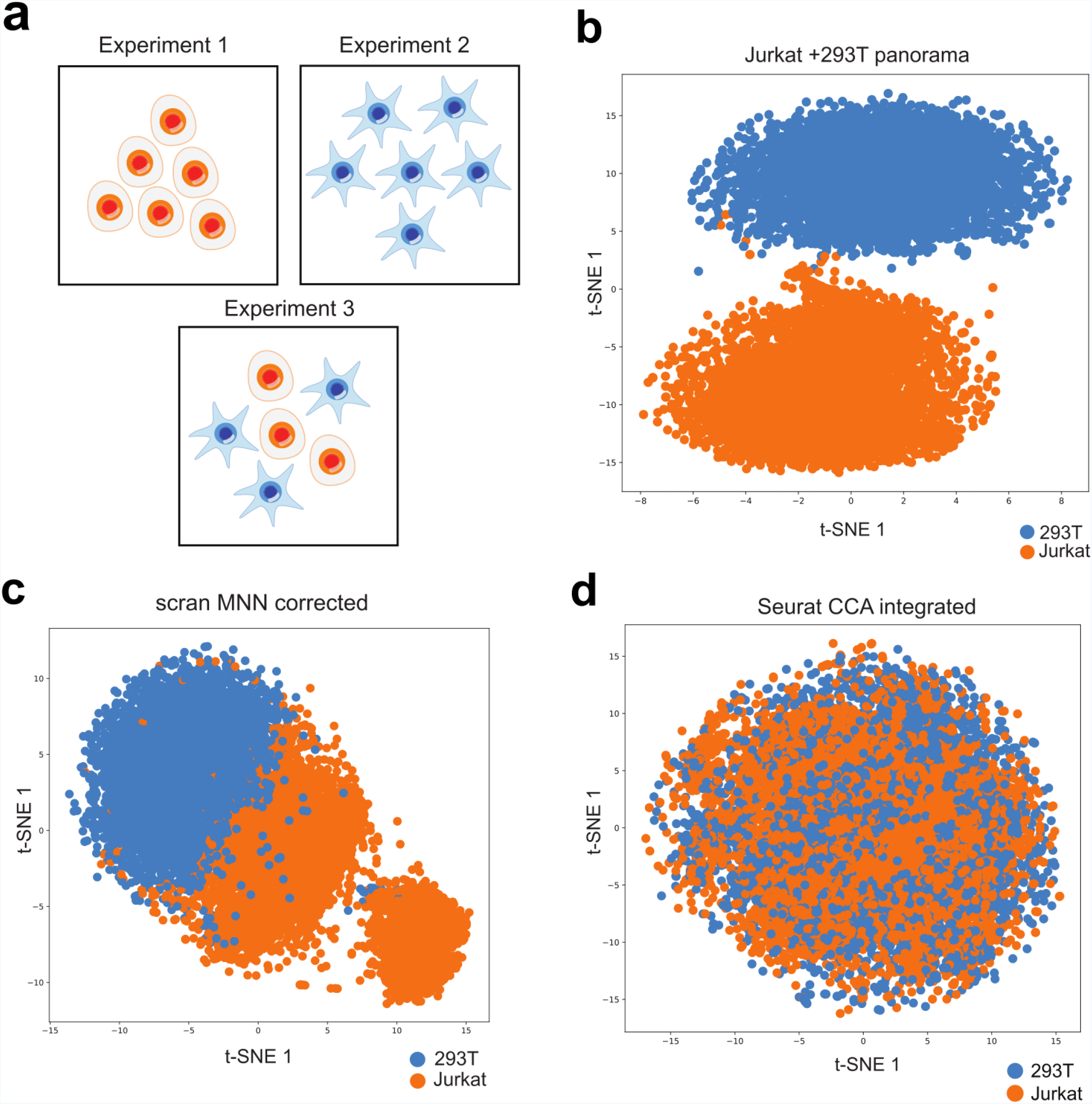
Scanorama correctly integrates a simple collection of data sets where other methods fail. (**a**) We apply Scanorama to a collection of three data sets^13^: one entirely of Jurkat cells (Experiment 1), one entirely of 293T cells (Experiment 2), and a 50:50 mixture of Jurkat and 293T cells (Experiment 3). (**b**) Our method correctly identifies Jurkat cells (orange) and 293T cells (blue) as two separate clusters. (**c,d**) Existing methods for scRNA-seq data set integration incorrectly merge a Jurkat data set and a 293T data set together first before subsequently incorporating a 293T/Jurkat mixture, forming clusters that do not correspond to actual cell types.

We then sought to demonstrate the ability of Scanorama to assemble a larger and more diverse set of cell types. In total, we ran our pipeline on 26 scRNA-seq data sets representing 9 different technologies and containing a total of 105,476 cells (**Fig. 3a, Supplementary Table 1**), each data set coming from a different scRNA-seq experiment. Scanorama correctly identifies data sets with the same cell types and merges them together such that they cluster by cell type instead of by experimental batch (**Fig. 3a-c**). Importantly, in contrast with existing methods, our algorithm does not merge disparate cell types together (**Fig. 3b,c; Supplementary Figure 3**), and correctly identifies a data set of mouse neurons as distinct from the cell types of all other data sets (**Fig. 3a**). One of the panoramas identified by Scanorama consists of two data sets of hematopoietic stem cells (HSCs)^14,15^ which, once corrected for batch effects and plotted along the first two principal components, reconstruct the expected HSC differentiation hierarchy (**Fig. 3d, Supplementary Fig. 4a,b**). We also observe cell type-specific clusters within panoramas of pancreatic islet cells (**Fig. 3e, Supplementary Fig. 5,6**) and peripheral blood mononuclear cells (**Supplementary Fig. 7,8**) but now have greater power to detect rare cell populations. For example, in the pancreatic islet panorama, we observe a cluster of cells consistent with a previously reported rare subpopulation of pancreatic beta cells marked by increased expression of endoplasmic reticulum (ER) stress genes (**Fig. 3e, Supplementary Fig. 6g,h**).^7^

**Figure 3.**
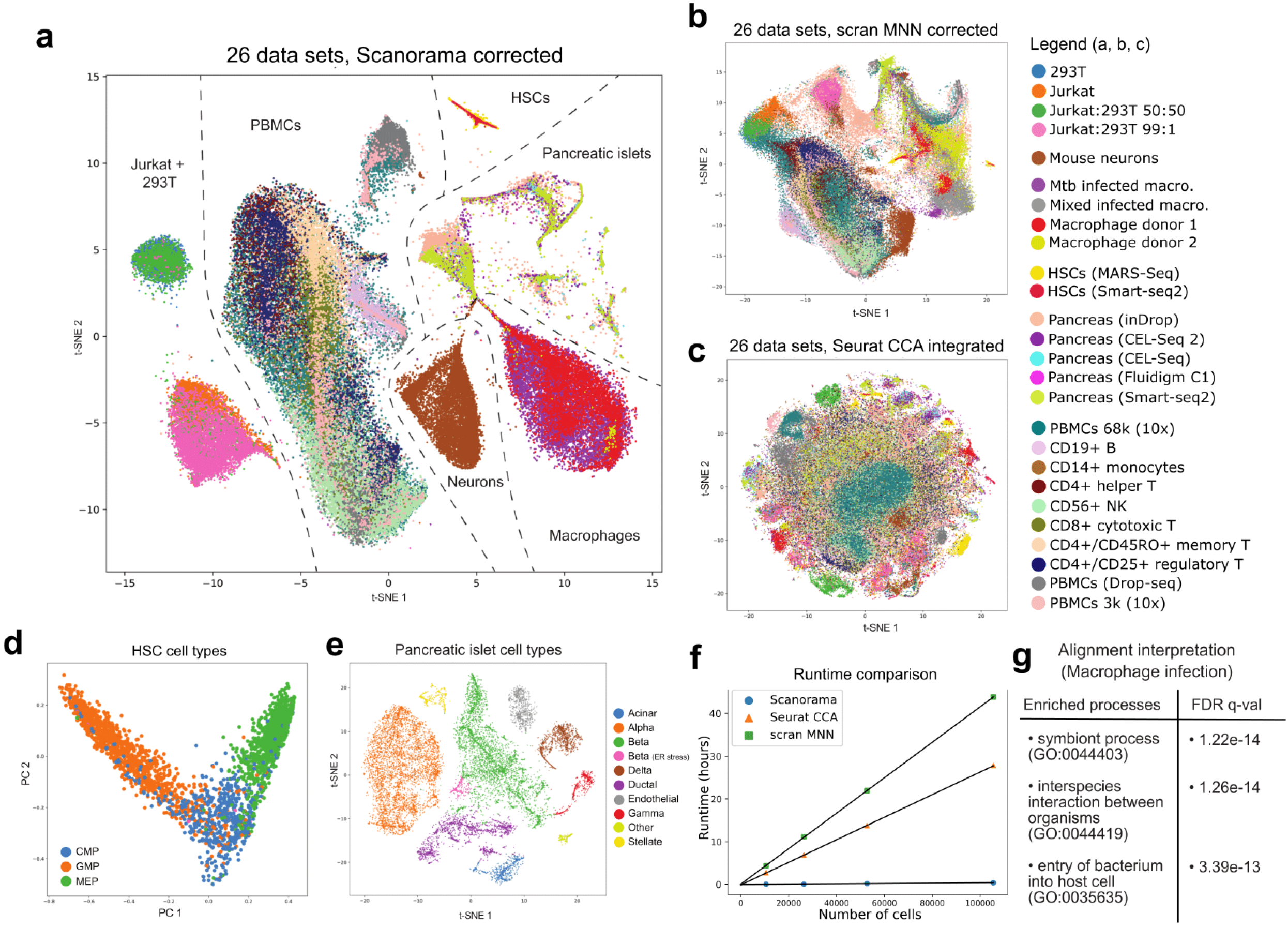
Panoramic integration of 26 single cell data sets across 9 different technologies. (**a**) t-SNE visualization of 109,618 cells after batch-correction by our method, with cells clustering by cell type instead of by batch (median Silhouette Coefficient of 0.34). (**b-c**) Other methods for scRNA-seq data set integration (scran MNN^6^ and Seurat CCA^7^) are not designed for heterogeneous data set integration and therefore naively merge all data sets into a single large cluster (median Silhouette Coefficient of 0.01 for scran MNN and −0.20 for Seurat CCA; **Supplementary Fig. 10**). (**d**) Scanorama automatically identifies a panorama of two hematopoietic stem cell (HSC) data sets^14,15^ and removes any significant difference due to experimental batch (likelihood-ratio = 2.7e-902; **Methods**). Visualizing the corrected HSCs along the first two principal components captures the “pseudo-temporal” arrangement of the cell types, in which granulocyte-macrophage progenitors (GMP) and megakaryocyte-erythrocytes (MEP) are derived from common myeloid progenitors (CMP). (**e**) Within a panorama of pancreatic islet cells from five different data sets^22–26^, we can observe clusters of distinct cellular subtypes including a previously reported rare subpopulation of beta cells marked by ER stress gene upregulation (**Supplementary Fig. 8g,h**). (**f**) Scanorama is significantly faster than current methods for scRNA-seq integration. (**g**) Interpreting the alignment between Mtb infected and uninfected macrophages reveals genes significantly enriched for processes related to bacterial infection.

Due to our algorithmic optimizations, Scanorama is also significantly faster than existing methods for scRNA-seq data set integration. In particular, to integrate our collection of 26 data sets containing 105,476 cells, Scanorama completes the nearest neighbor search to find panoramas in roughly five minutes and performs batch-correction of all panoramas in under 25 minutes. In contrast, existing methods require more than 27 hours to integrate the same collection of data sets (**Fig. 3f**) using about the same amount of memory (**Supplementary Fig. 9**) while also performing poorly at preserving real biological heterogeneity in the integrated result (**Fig. 3b,c, Supplementary Fig. 10**).

The alignments made by Scanorama are also interpretable, allowing researchers to examine the genes that significantly contribute to the alignment between two data sets and better understand the technical or biological origin of discrepancies between experiments. For example, Scanorama aligned two data sets consisting of the same cell type but separated by a known biological difference, i.e., two macrophage data sets with and without *Mycobacterium tuberculosis* (Mtb) infection. We found that the alignment of macrophage populations between these data sets is directed by genes that are enriched for processes relating to bacterial infection (**Fig. 3g, Supplementary Table 2**), indicating that our alignment vectors can be examined for biological significance. Such information gives researchers a better understanding of the alignment between data sets, which may optionally be corrected by our pipeline.

As researchers work to assemble a more complete picture of diverse biological functions at a single-cell resolution, the need to integrate heterogeneous experiments also increases. Our algorithm provides a robust and efficient solution to this problem, which we make publicly available at http://scanorama.csail.mit.edu. We note that Scanorama is modular and can be used alongside other tools, which includes assembling the reference data set required for projective methods^16^ or combined with different methods for scRNA-seq clustering and visualization.^17–19^ While we currently align all high-quality cells, random or structure-preserving sampling of the data^20^ could further improve computational efficiency. Our algorithm may also be used to interrogate a system in which the underlying cell states are unknown and are not guaranteed to be shared among different experiments, which may elucidate novel biological patterns. Scanorama provides a robust platform for integrating complex biological data sets acquired using scRNA-seq, thereby enabling the identification of biological paradigms, across multiple diseases, tissues, and biological conditions, independent of sequencing technology or cell subpopulation frequency.

## Methods

### Data sets and data processing

We used the following publicly-available data sets:

- 293T cells from Zheng *et al*. (2017)^13^ https://support.10xgenomics.com/single-cell-gene-expression/datasets/1.1.0/293t)
- Jurkat cells from Zheng *et al.* (2017)13 (https://support.10xgenomics.com/single-cell-gene-expression/datasets/1.1.0/jurkat)
- 50:50 Jurkat:293T cell mixture from Zheng *et al.* (2017)^13^ (https://support.10xgenomics.com/single-cell-gene-expression/datasets/1.1.0/jurkat:293t_50:50)
- 99:1 Jurkat:293T cell mixture from Zheng *et al.* (2017)^13^
- (https://support.10xgenomics.com/single-cell-gene-expression/datasets/1.1.0/jurkat_293t_99_1)
- Mouse neurons from 10x Genomics (unpublished)
- (https://support.10xgenomics.com/single-cell-gene-expression/datasets/2.1.0/neuron_9k)
- Macrophages (Mtb infected) (pending final GEO approval)
- Macrophages (Mtb exposed) from Gierahn *et al.* (2017)^21^ (GSE92495)
- Macrophages (unexposed) (pending final GEO approval)
- Macrophages (unexposed) from Gierahn *et al*. (2017)^21^ (GSE92495)
- Mouse hematopoietic stem cells (HSCs) from Paul *et al*. (2015)^14^ (GSE72857)
- Mouse HSCs from Nestorowa *et al*. (2016)^15^ (GSE81682)
- Human pancreatic islet cells from Baron *et al.* (2016)^22^ (GSE84133)
- Human pancreatic islet cells from Muraro *et al*. (2016)^23^ (GSE85241)
- Human pancreatic islet cells from Grün *et al.* (2016)^24^ (GSE81076)
- Human pancreatic islet cells from Lawlor *et al.* (2017)^25^ (GSE86469)
- Human pancreatic islet cells from Segerstolpe *et al.* (2016)^26^ (E-MTAB-5061)
- Human PBMCs from Zheng *et al.* (2017)^13^ (https://support.10xgenomics.com/single-cell-gene-expression/datasets/1.1.0/fresh_68k_pbmc_donor_a)
- Human CD19+ B cells from Zheng *et al.* (2017)^13^
- (https://support.10xgenomics.com/single-cell-gene-expression/datasets/1.1.0/b_cells)
- Human CD14+ monocytes from Zheng *et al*. (2017)^13^
- (https://support.10xgenomics.com/single-cell-gene-expression/datasets/1.1.0/cd14_monocytes)
- Human CD4+ helper T cells from Zheng *et al*. (2017)^13^
- (https://support.10xgenomics.com/single-cell-gene-expression/datasets/1.1.0/cd4_t_helper)
- Human CD56+ natural killer cells from Zheng *et al*. (2017)^13^
- (https://support.10xgenomics.com/single-cell-gene-expression/datasets/1.1.0/cd56_nk)
- Human CD8+ cytotoxic T cells from Zheng *et al*. (2017)^13^
- (https://support.10xgenomics.com/single-cell-gene-expression/datasets/1.1.0/cytotoxic_t)
- Human CD4+/CD45RO+ memory T cells from Zheng *et al*. (2017)^13^
- (https://support.10xgenomics.com/single-cell-gene-expression/datasets/1.1.0/memory_t)
- Human CD4+/CD25+ regulatory T cells from Zheng *et al.* (2017)^13^
- (https://support.10xgenomics.com/single-cell-gene-expression/datasets/1.1.0/regulatory_t)
- Human PBMCs from Kang *et al.* (2018)^27^ (GSE96583)
- Human PBMCs from 10x Genomics (unpublished)
- (https://support.10xgenomics.com/single-cell-gene-expression/datasets/1.1.0/pbmc3k)

In each data set, we removed low-quality cells by including only those with at least 600 identified genes. When searching for scRNA-seq panoramas, we only consider the genes that are present in all data sets and *l*_2_-normalize the expression values for each cell for scale-invariant comparison. In our study, there were 5,216 genes present across all 26 data sets, each data set containing between 90 and 18,018 cells, and which in total contained 105,476 high-quality cells (**Supplementary Table 1**).

### Dimensionality reduction using randomized SVD

We compute a compressed, low-dimensional embedding of the gene expression values for each cell by taking the SVD of the combined cell-by-gene expression matrix, taking inspiration from different compressive techniques for other biological problems.^28^ The SVD is normally very expensive to compute on large matrices, so we leverage an efficient, randomized approach to find an approximate SVD^10^, which is implemented in the fbpca Python package (http://fbpca.readthedocs.io/en/latest/). We use a reduced dimension of 100 in all of our experiments.

### All-to-all data set matching

For each data set, we query for its cells’ nearest neighbors among the cells of all remaining data sets in the low-dimensional embedding space. After repeating this for all data sets, we find all instances where a cell in one data set is the nearest neighbor of a cell in another data set, and vice versa. Additional algorithm descriptions are given in the **Supplementary Materials**.

### Approximate nearest neighbors using locality sensitive hashing

To greatly accelerate our nearest neighbor queries, our algorithm conducts an approximate search based on locality sensitive hashing, where multiple trees of random hyperplanes, used as hash functions, divide the search space of the points in the query set.^11,12^ We use the Annoy C++/Python package (https://github.com/spotify/annoy), a memory efficient implementation of this algorithm.

### Nonlinear data set merging and panorama stitching

Once mutual nearest neighbor matches are identified, we merge data sets together into larger panoramas. We build upon the nonlinear batch-correction strategy of Haghverdi *et al.*^6^ that maps one scRNA-seq data set onto another by computing translation vectors in the full gene expression space for all cells in a data set. Translation vectors for each cell are obtained as a weighted average of the matching vectors (defined by the pairs of matched cells), where a Gaussian kernel function upweights matching vectors belonging to nearby points. We order pairs of data sets based on the percentages of cells in the data sets that are involved in a matching and use this ordering to build panoramas of data sets by successively merging a data set into a panorama or using the pair of data sets to merge two panoramas together using the nonlinear correction procedure described above. Additional algorithm descriptions are given in the **Supplementary Materials**.

### Interpreting alignments

Since aligning and merging two data sets is based on a set of matching vectors, we reasoned that the magnitude of these vectors along the dimensions of the gene expression space would allow us to identify genes with a significant contribution to the direction of the alignment. To interpret an alignment between two data sets, we averaged the matching vectors across all matchings and compared the absolute magnitude of each of the values in the mean vector to the corresponding absolute magnitudes of a “null set” of vectors. We constructed a null distribution by randomly pairing cells between the two data sets and computing the vectors obtained from each pair (excluding pairs in the matching itself). We used a null distribution of 100,000 vectors to calculate a permutation-based p-value for each of the genes. Genes were put into a target list at a false discovery rate (FDR)^29^ of less than 0.05, for which we looked for gene ontology (GO) process enrichment among the background set of 5,216 genes using the GOrilla web tool (http://cbl-gorilla.cs.technion.ac.il/).^30^

### t-SNE visualization

We modified the implementation of t-Distributed Stochastic Neighbor Embedding (t-SNE)^31^ in scikit-learn^32^ by replacing the exact nearest neighbors search phase with an approximate nearest neighbors search using the same locality sensitive hashing algorithm and implementation as in our data set matching procedure. This modification was done to improve the runtime of the default scikit-learn t-SNE when visualizing our results and is included in the code package for our algorithm.

### Clustering performance

Previous scRNA-seq analyses^6,33^ have used the Silhouette Coefficient^34^ as a quantitative measure of clustering performance. The Silhouette Coefficient is calculated using the mean of the distances from cell *i* to all other cells of the same type (*a_i_*) and the mean of the distances from cell *i* to all other cells that belong to the cell type that is nearest to the cell type of *i* (*b_i_*). The Silhouette Coefficient for a cell is 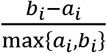, taking values between 1 and -1, inclusive, where higher values indicating better clustering performance. Intuitively, the Silhouette Coefficient improves if a cell is close to other cells of the same type and far from cells of a different type. We computed the Silhouette Coefficient using the t-SNE embeddings of all cells across our collection of 26 data sets integrated by Scanorama, scran MNN, and Seurat CCA (**Supplementary Fig. 10**). We use the Silhouette Coefficient implementation provided by scikit-learn.

### Simulation of non-overlapping data sets

We simulated data sets by sampling from a 2-dimensional Gaussian mixture model with four components and isotropic noise. We then projected all of our data sets into a 100-dimensional space using a matrix with Gaussian noise entries. We also simulated batch effects for each data set by adding a vector of Gaussian noise to all entries in a data set, using a different random vector (i.e., a different batch effect) for each data set.

### Panorama of 293T and Jurkat cells

We obtained three separate data sets consisting of 293T cells, Jurkat cells, and a 50:50 mixture of 293T and Jurkat cells from 10x Genomics.^13^ These data sets were processed, aligned, and merged using the procedure described previously to give a total of 9,530 cells. We defined cell types using labels from the original study.

### Panorama of hematopoietic stem cells (HSCs)

Two publicly available data sets^14,15^ of HSCs were processed, aligned, and merged using the procedure described previously to give a total of 3,175 cells. We used the cell types that were reported by both studies, and we examined the expression of marker genes indicating erythropoiesis for additional validation (**Supplementary Fig. 4c-e**). We quantified the quality of our batch correction by computing the likelihood-ratio using the likelihood that the corrected MARS-Seq data set came from the same distribution as the uncorrected MARS-Seq data set (*H*_0_) or from the same distribution as the corrected Smart-seq 2 data set (*H*_1_), where we can more confidently reject the null hypothesis *H*_0_ if the likelihood-ratio 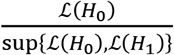 is very small. We modeled each distribution with a three-component Gaussian mixture model.

### Panorama of pancreatic islets

Five publicly-available pancreatic islet data sets^22–26^ were processed, aligned, and merged using the procedure described previously to give a total of 15,921 cells (**Supplementary Fig. 5**). We k-means clustered the cells in the corrected gene expression space, obtaining 40 clusters, and assigned cell types to each cluster based on cell type labels provided by previous clustering analyses^7,23,25^ and the relative expression levels of cell type-specific marker genes (**Supplementary Fig. 6**). This allowed us to identify a cluster corresponding to a rare subpopulation of beta cells with upregulated ER stress genes, which we identified using previously inferred labels^7^ for one of the data sets^22^ and by confirming upregulation of the marker genes *HERPUD1* and *GADD45A* in cells from all data sets within that cluster (**Supplementary Fig. 6g,h**).

### Panorama of peripheral blood mononuclear cells (PBMCs)

Ten publicly-available data sets^13,27^ involving PBMCs, or cell types found in PBMCs, were processed, aligned, and merged using the procedure described previously to give a total of 47,994 cells. Cell types were either experimentally determined^13^ using fluorescence activated cell sorting (FACS) or inferred by previous clustering analyses^6,7^ (**Supplementary Fig. 7**), and we examined the expression levels of cell type-specific marker genes for additional validation (**Supplementary Fig. 8**).

### Macrophage alignment interpretation

Four publicly available data sets^21^ of macrophages, sorted by FACS for Mtb infection, were processed and aligned using the procedure described previously to give a total of 15,337 cells. Once the matching vectors between the Mtb infected and uninfected data sets were determined, we used the procedure described previously to identify genes and functional processes that significantly contribute to the discrepancy between the two data sets (**Supplementary Table 2**).

### Runtime and memory profiling

We used Python’s time module to obtain runtime measurements for the alignment and merging portions of our algorithm and used the top program in Linux (Ubuntu 17.04) to make periodic memory measurements. We also randomly subsampled sets of 10,547 (10%), 26,369 (25%), and 52,738 (50%) cells from our total of 105,476 cells and measured the runtime and memory of our algorithm on the subsampled data. We compared computational resource usage to two methods, Seurat CCA^7^ (with 15 canonical correlation vectors) and scran MNN^6^, using their default parameters. For a fair comparison, we used the same preprocessed data and only measured the resources required for the portions of the methods responsible for alignment and data set integration. We used R’s proc.time function and Linux’s top to measure runtime and memory usage, respectively, of these programs. All methods were limited to 10 cores and run on a 2.30 GHz Intel Xeon E5-2650v3 CPU with 384 GB of RAM.

### Data and code availability

The data and code used in this study are available at http://scanorama.csail.mit.edu.

## Acknowledgements

B. Hie is partially supported by NIH grant R01GM081871 (to B. Berger). We thank H. Cho, S. Nyquist, and L. Schaeffer for valuable discussions and feedback. We thank S. Tovmasian for assistance in preparing the manuscript.

## Author Contributions

B. Hie developed the algorithm, built the computational tools, and performed the analysis in discussions with B. Bryson and B. Berger. B. Berger supervised the research. All authors wrote the manuscript.

## Competing Interests

The authors declare no competing interests.

